# Automated movement assessment in stroke rehabilitation

**DOI:** 10.1101/2021.06.25.449936

**Authors:** Tamim Ahmed, Kowshik Thopalli, Thanassis Rikakis, Pavan Turaga, Aisling Kelliher, Jia-Bin Huang, Steve Wolf

**Author notes:** Correspondence: Corresponding Author, Tamim Ahmed, Interactive Neurorehabilitation Lab, Department of Biomedical Engineering, Virginia, Tech, Blacksburg, VA, 24061, USA, Kowshik Thopalli, Geometric Media Lab, Department of Electrical and Computer Engineering, Arizona, State University, Tempe, Arizona, 85281, USA.

## Abstract

We are developing a system for long term Semi-Automated Rehabilitation At the Home (SARAH) that relies on low-cost and unobtrusive video-based sensing. We present a cyber-human methodology used by the SARAH system for automated assessment of upper extremity stroke rehabilitation at the home. We propose a hierarchical model for automatically segmenting stroke survivor’s movements and generating training task performance assessment scores during rehabilitation. The hierarchical model fuses expert therapist knowledge-based approaches with data-driven techniques. The expert knowledge is more observable in the higher layers of the hierarchy (task and segment) and therefore more accessible to algorithms incorporating high level constraints relating to activity structure (i.e. type and order of segments per task). We utilize an HMM and a Decision Tree model to connect these high level priors to data driven analysis. The lower layers (RGB images and raw kinematics) need to be addressed primarily through data driven techniques. We use a transformer based architecture operating on low-level action features (tracking of individual body joints and objects) and a Multi-Stage Temporal Convolutional Network(MS-TCN) operating on raw RGB images. We develop a sequence combining these complimentary algorithms effectively, thus encoding the information from different layers of the movement hierarchy. Through this combination, we produce a robust segmentation and task assessment results on noisy, variable and limited data, which is characteristic of low cost video capture of rehabilitation at the home. Our proposed approach achieves 85% accuracy in per-frame labeling, 99% accuracy in segment classification and 93% accuracy in task completion assessment. Although the methodology proposed in this paper applies to upper extremity rehabilitation using the SARAH system, it can potentially be used, with minor alterations, to assist automation in many other movement rehabilitation contexts (i.e. lower extremity training for neurological accidents).

## 1 Introduction

As the US and global populations age, we observe an increasing need for effective and accessible rehabilitation services for survivable debilitating illnesses and injuries, such as stroke and degenerative arthritis (Haghi M, 2017; Pantelopoulos and Bourbakis, 2010). Effective rehabilitation requires intensive training and the ability to adapt the training program based on patient progress and therapeutic judgment (Kleim and Jones, 2008). Intensive and adaptive rehabilitation is challenging to administer in an accessible and affordable way; high intensity therapy necessitates frequent trips to the clinic (usually supported by a caregiver), and significant one-on-one time with rehabilitation experts (Lang et al., 2013). Adaptation requires a standardized, evidence-based approach, coordinated amongst many specialists (Chen et al., 2011b; Duff et al., 2008; Lehrer et al., 2011). Active participation by the patient is also critical for improving self-efficacy and program adherence (Picha and Howell, 2017), although, without significant dedicated effort from a caregiver, in many cases, active participation and adherence are difficult to achieve (Donelan et al., 2002).

Telemedicine and telehealth are gaining significance as viable approaches for delivering health and wellness at the home and in the community at scale (Clinic, 2015). Applying existing telemedicine approaches to physical rehabilitation in the home is not yet possible, owing to the challenges of automating the observation, assessment, and therapy adaptation process used by expert therapists. For upper extremity rehabilitation for stroke survivors, which is the focus of this paper, more than 30 low-level movement features need to be tracked as the patient performs functional tasks in order to precisely and quantitatively characterize movement impairment (Chen et al., 2011b). High precision sensing and tracking systems can work well in spacious and supervised clinical environments, but are currently not yet appropriate for a typical home setting. The use of marker-based tracking systems or complex exoskeletons are simply too expensive, challenging to use, and obtrusive in the home (Deaver et al., 2019; Hanley et al., 2018; Reinkensmeyer et al., 2017).

Banks of video camera arrays can seem intrusive in the home and lead patients and/or their families to feel as if they are under surveillance (Duff et al., 2008). More promisingly, networks of wearable technologies (e.g. IMUs, smart skins, pressure sensors) can provide useful tracking data for overall movement and detailed features, but they can also be hard to put on correctly, irritating to wear for long periods of time, and sometimes require a perceived excessive number of wearables to capture all movement features correctly (Freedson et al., 1998). A final concern with respect to the patient’s home environment concerns the physical footprint of any technology introduced into their home. Disturbing the home setting can be understood negatively, which has the knock-on effect of reducing adoption by stroke survivors and/or other family members in the home (Axelrod et al., 2009).

That said, accurate, low-cost capture of movement data is only part of the challenge. Automation of assessment is also difficult because the processes used by therapists are largely tacit and not well standardized (Levin et al., 2008). Clinicians are trained to use validated clinical measures (e.g. Fugl-Meyer, ARAT, and WMFT), utilizing a small range of quantitative scales (0-2, 0-3, and 0-5 respectively) for assessing performance of functional tasks that map to activities of daily life (Rabadi and Rabadi, 2006; Wolf et al., 2001). These scales provide standardized rubrics for realizing this assessment and use a uniform activity space with exact measurements for each subcomponent of the space to assist the standardized performance of the tasks. The high-level rating of task performance provided by experts through these measures is difficult to map directly to specific aspects of movement and related detailed kinematics extracted through computational means. Even expert clinicians cannot simultaneously observe all aspects of upper extremity pathological movement or compare such observations to a standardized value. There is considerable evidence showing that individual therapists direct their attention towards different elements and assess them differently when evaluating performance in situ and real-time, or when later rating videos of performance (Chen et al., 2011a; Cirstea and Levin, 2007; Nordin et al., 2014; Wolf et al., 2006). The relation of movement quality to function is therefore difficult to ascertain in a standardized quantitative manner (Levin et al., 2008; Chen et al., 2011b). This results in approaches for structuring and customizing therapy that are partly based on subjective experience rather than a standardized quantifiable framework (Cirstea and Levin, 2007; Levin et al., 2008). In turn, this results in a lack of large-scale data on the structuring and customization of adaptive therapy, and on the effects of customization and adaptation choices on functionality in everyday life (Reinkensmeyer et al., 2016). Therefore, full scale automation of the real-time functions of the therapist at the home is not yet feasible.

In light of these limitations, we are developing the novel Semi-Automated Rehabilitation At the Home (SARAH) system. The SARAH system comprises two video cameras, a tablet computer, a flexible activity mat, and eight custom-designed 3D printed objects, as shown in Fig. 2 below. The objects are designed to support a broad range of perceived affordances (Norman, 2002), meaning they can be gripped, moved, and manipulated in a wide variety of ways (Kelliher et al., 2019). Each object is unique in terms of size and color to assist identification of objects by patients, and to enable easier identification and tracking using computer vision methods. The flexible activity mat is screen-printed with high-contrast guidance lines indicating to the patient the four primary activity spaces (near and distal ipsilateral and contralateral) through increasingly colored lines. In addition, four rows of circles on the mat assist the computer vision system with boundary detection between activity space for consistently analyzing patient activities.

The system can be easily installed on a regular kitchen or living room Table. The two cameras initiate and record only during training, and they are activated and controlled by the patient using a custom-designed application on the tablet computer. The system aims to integrate expert knowledge with data driven algorithms to realize coarse real-time automated assessment of movement of stroke survivors during therapy at the home (Kelliher et al., 2020). This assessment can drive high-level feedback on results and performance after execution of each training task, to assist patients with self assessment, and to help them plan their next attempt(s) of the training task. Daily summaries of the interactive training will be transmitted to remote therapists to assess overall progress (within and across sessions), adjust the therapy structure, and provide text or audio based feedback and directions to the patient via the tablet. Continuous and effective training monitoring accompanied by feedback on the patient’s immediate performance, combined with expert customization of therapy to their needs and learning styles, increases the likelihood of patients adopting home-based rehabilitation systems (Picha and Howell, 2017).

This paper focuses on our hybrid knowledge-based data-driven approach to automated assessment of human movement in the home. Our approach leverages expert rubrics for standardized rating of overall task performance to inform automated rating of movement performance based on low cost, limited, noisy, and variable kinematic data. The assessment process and outcomes need to be compatible with therapist assessment approaches so as to assist remote therapists in using summaries of the computational assessment when remotely monitoring progress and structuring therapy at the home.

Our approach has two components: i) making the expert raters process as observable as possible; and ii) leveraging the expert rating process to inform the structuring and improved performance of computational algorithms. In previous publications, we have presented in detail our research and development activities for the first component (Rikakis et al., 2018; Clark and Kelliher, 2021; Kelliher et al., 2020). Inspired by clinical measures for rating rehabilitation movement, we developed the SARAH system to utilize a standardized activity space with eight well defined sub-spaces that are drawn as bounding boxes on the video capture of therapy (see Fig. 6). We designed the SARAH training objects to facilitate generalized mapping of training tasks to ADLs for the patients, while also facilitating tracking through low-cost video cameras (Kelliher et al., 2019). We used participatory design processes and custom-designed interactive video rating tools to help expert therapists reveal and reflect on their rating process and internalized (tacit) rating schemas (Kelliher et al., 2020).

In order to manage the complexity of real-time movement observation and to make generalizable observations across different therapy tasks, therapists tend to segment tasks into a few segments that can be combined in different sequences to generate targeted therapy tasks. Even though most therapists use intuitive segmentation of movement for observation and assessment, the segment vocabulary is *not* standardized. We worked with expert therapists to standardize the segment vocabulary into a state machine that can produce all 15 tasks of the SARAH system (Kelliher et al., 2020). The segments are: Initiation + Progression + Termination (IPT), Manipulate and Transport (MTR), Complex Manipulation and Transport (CMTR), and Release and Return (RR). As an example, a drinking related task can be described by the following codification: subject reaches out and grasps a cone object (IPT) and brings it to their mouth (MTR), then returns the object to the original position (MTR), and releases the object and returns the hand to the rest position (RR).

To make the assessment of segments in real-time manageable, the therapist significantly limits the features observed per segment. This limitation is achieved by using their own experience to develop a probabilistic filtering of irrelevant low-level features for a segment (i.e. digit positioning is likely not that relevant to movement initiation), and probabilistic composite observation of relevant features (i.e. a strategy for quick impressions of shoulder and torso compensation during movement initiation). This process is not well standardized as the filtering and compositing activities are based on individual experience and training. We further worked with expert therapists to define a consensus-limited set of composite movement features that are important when assessing the performance of each segment in our model Kelliher et al. (2020). For example, the resulting rubric identifies four key features to assess during the Complex Manipulation and Transport stage: i) appropriate initial finger positioning, ii) appropriate finger motion after positioning, iii) appropriate limb motion following finger positioning, and iv) limb trajectory with appropriate accuracy. The rubric also establishes operational definitions of terms used to evaluate movement quality and inform rating. For example, the word “appropriate” used in the above instructions is defined as “the range, direction, and timing of the movement component for the task compared to that expected for the less impaired upper extremity.” Although therapists do not explicitly track and assess raw kinematic features, in previous work we proposed computational approaches connecting therapist’s assessment of composite features to computationally tracked raw kinematics (Venkataraman et al., 2014; Chen et al., 2011b).

## 2 MATERIALS AND METHODS

### 2.1 Analysis Framework

Based on the background discussed above, we propose a cyber-human movement assessment hierarchy for upper extremity rehabilitation. The five layers of the hierarchy (listed top to bottom) are: overall task rating, segment rating, composite feature assessment, raw kinematics, and raw RGB images (see also Fig. 1). As we move down the hierarchy, the expert knowledge becomes less observable and less standardized. Computational knowledge works in reverse, with more confidence in the lower levels (raw kinematics or derivatives) that gradually diminishes moving up in the hierarchy towards complex decision-making relying on expert experience (e.g., overall task rating). The proposed hierarchy aims to reveal the grammar (or compositional structure) of therapy, from the signal level to the meaningful activity level (training tasks). Knowing the composition rules of a complex human activity, whether that is language, sport, or a board game like chess, facilitates the meaningful computational analysis of the activity in a manner that is comprehended and leveraged by expert trainers and trainees Simon (1981); Venkataraman et al. (2016). Although the hierarchy proposed in this paper is established for upper extremity rehabilitation using the SARAH system, it can potentially be used, with minor alterations, to assist automation in many other movement rehabilitation contexts (i.e. lower extremity training for neurological accidents). Our proposed methodology can also transfer to other complex human activity contexts (i.e. training firefighters or athletes).

**Figure 1.**
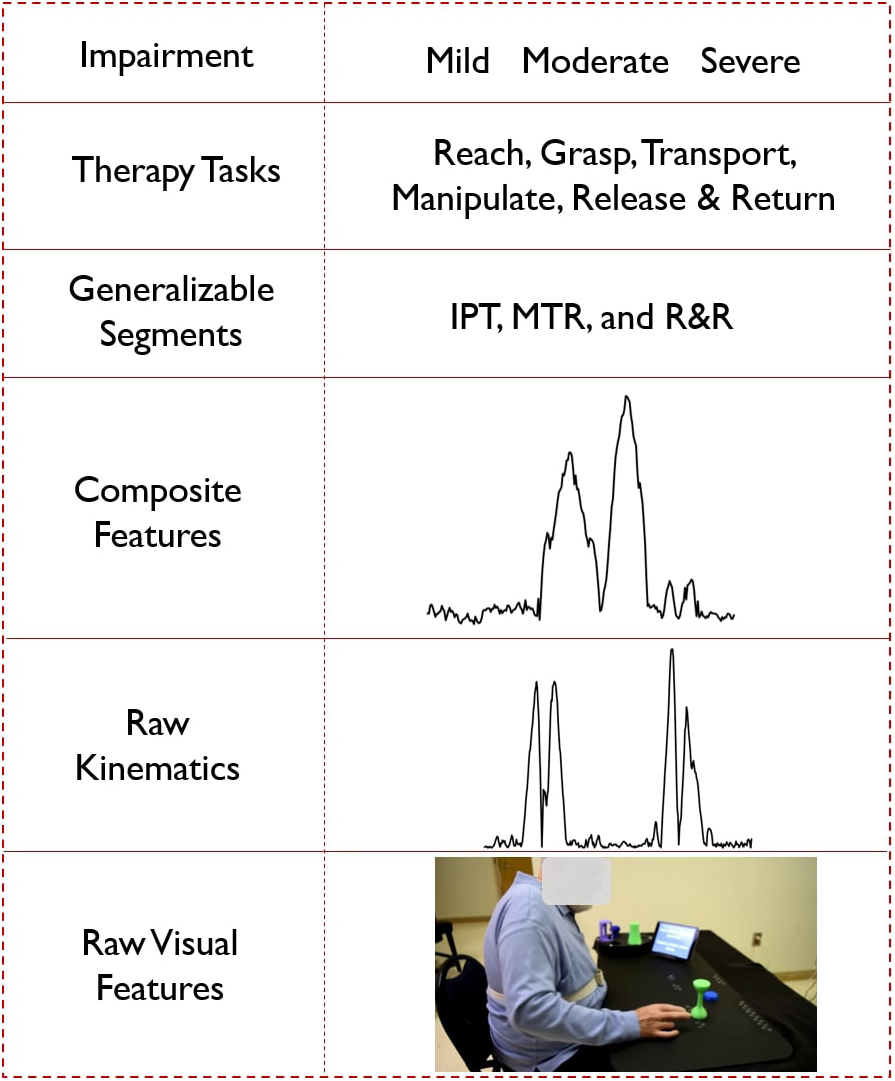
Hierarchical representation of the cyber-human system; (from the bottom) a video frame as an example of raw visual features, velocity profile as an example of raw kinematic features, composite feature for task 5 generated using PCA, segment labels, different types of tasks and impairment levels

**Figure 2.**
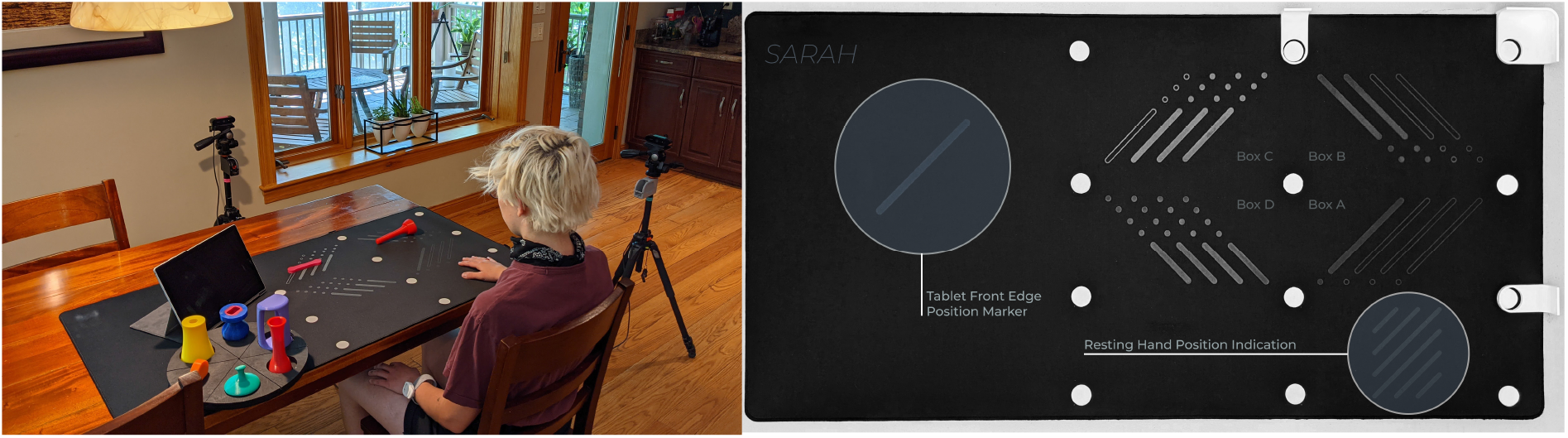
(a) SARAH system and objects setup; (b) SARAH activity mat.

In the following sections, we show how we leverage the observable expert knowledge of the higher levels of the hierarchy to improve the performance of computational algorithms using raw kinematics and visual features for automated task segmentation, segment classification, and task performance assessment at the home. As the patient selects the task they want to attempt using the tablet computer, the system knows the desired sequence of segments for satisfactory performance of the task. The expected topology of each segment of each task within our standardized activity space is also specified. We can thus utilize a feature-matching approach to measure the ‘distance’ between the observed sequence of segments by the system and the expected sequence of states for the performed task to determine whether the task was successfully performed.

The state-time characteristics of our hierarchy (i.e segment state machine) motivates our use of automated segmentation though a hidden Markov model (HMM). We also leverage recent advances in machine-learning models for processing video data. Specifically, we tested the use of transformers (Vaswani et al., 2017) operating on low-level action features (tracking of individual body joints and object tracking). We also tested machine learning algorithms that operate on raw RGB images (MS-TCN) (Farha and Gall, 2019). We chose the MS-TCN algorithm because its classifiers are trained on recognizing human activity based on kinetics and thus can synergize well with automated assessment of task performance in rehabilitation. All three types of algorithms required slightly different features for achieving best performance in connecting the raw data layers to the segment layer (and achieving automated segmentation). In addition, the features used by the algorithms were different from the composite features used by the human experts. This established the segments layer as the key integration layer for different assessment approaches, and reinforced the important role of a generalizable segment vocabulary proposed by expert therapists. Each approach showed different strengths and weaknesses, therefore we utilized an ensemble approach across the algorithms to finalize the automated segmentation and segment classification decisions. Automated assessment of therapy at the home using the SARAH system will need to be realized through noisy and variable data because of the low cost infrastructure, the varying environments of installation in the home, and the significant variation of movement impairment of stroke survivors (Rikakis et al., 2018). Furthermore, at the early stages of implementation of the SARAH system the data sets will be limited. To explore the resilience of our methodology we recreated these conditions in a clinic and gathered a limited data set for testing the algorithms. As we show in detail in the next section, even though the individual data layers of our cyber-human hierarchy are noisy, the composite automated decisions (successful performance of individual tasks and sequence of tasks assigned) produced across layers are robust. In future work, we will explore the expanded use of this approach for detailed automated assessment of elements of movement quality and the relation of movement quality to functionality.

### 2.2 Data Collection

We used an earlier version of the SARAH system (Kelliher et al., 2017) to record videos of nine stroke survivors (seven men and two women) performing the first 12 SARAH tasks. The nine stroke survivors had different levels of impairment ranging from mild/moderate (Fugl-Meyer score between 30-55) to moderate impairment (Fugl-Meyer score greater than 55) and had different types of movement challenges. Thus our overall dataset represents good range of conditions and movement variability. The participants in the study were asked to attempt each of the first 12 SARAH tasks, repeating each task four times if possible. The majority of patients could not perform the final two of the 12 SARAH tasks since the last two are the most difficult. For this reason, in our analysis below, we only use 450 videos of the individual performances of the first ten tasks by the nine patients.

The movement of the patients in this study was recorded using one consumer grade video camera (we used the side view camera of the SARAH system to capture the profile of the body and impaired arm as this is the preferred viewing point of the therapist). The camera placement instructions were relatively high-level to ensure that they could be implemented quickly and without interfering with the patient’s therapy session. The research assistant collecting the videos was asked to place the camera on the side of the impaired limb of the patient. The camera had to be far enough so as to not interfere with the performance of the tasks and be able to capture the full upper body of the patient. No specific distance, height or viewing angle were given for camera placement. As expected, this process provided minimal interference and could be realized very quickly but produced high variability in the captured videos in terms of location of the patient, activity space in the image, lighting, camera height, and viewing angle.

## 3 AUTOMATED SEGMENTATION AND SEGMENT CLASSIFICATION

Our segmentation and classification framework is illustrated in Fig. 3. It has three sets of blocks denoted with different colors in the figure. The first set of blocks are fine-tuned or pre-trained models that we implement as feature extractor. The second set of blocks include different algorithms such as HMM, Transformer, MSTCN++, and RBBDT. The third set of blocks are knowledge constrained formulas that we use as feature extractor for the RBBDT, HMM and also use to predict segment blocks and do task assessment. The HMM, Transformer, and MSTCN++ blocks generate per-frame state probabilities from the input video data. We generate the per-frame segment labels by a fusion of the state probabilities. With the incorporation of the design constrained denoising and the candidates from the RBBDT block (the decision tree), we calculate segment blocks from the per-frame ensemble predictions. Finally, we calculate task performance assessment scores using the segment blocks. The following subsections cover the detailed description of different blocks in the analysis framework.

**Figure 3.**
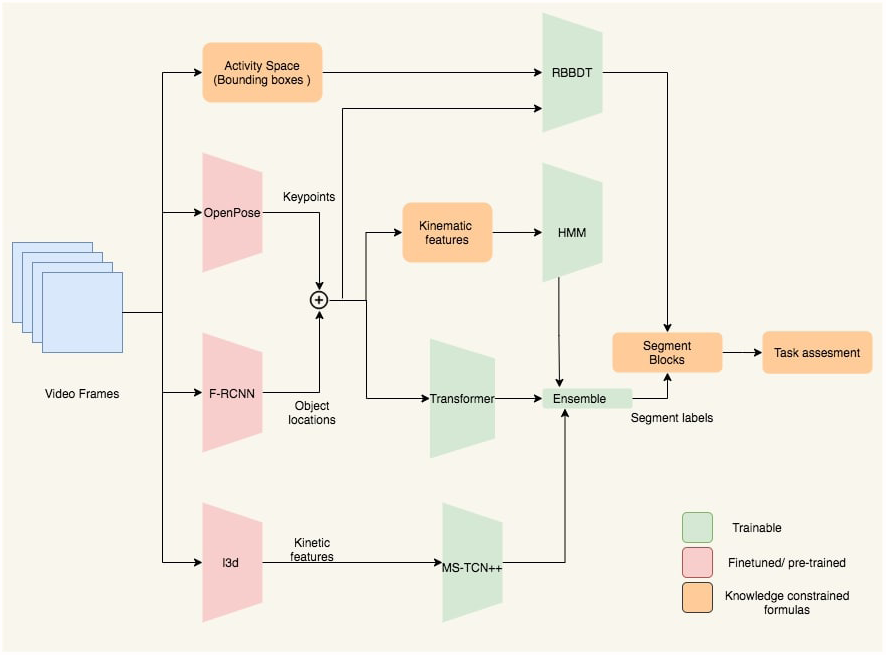
Block diagram of the proposed analysis framework. Keypoints and object locations are extracted from OpenPose and Faster-RCNN followed by a kalman filtering respectively. RBBDT along with the model ensemble determine the segment blocks from the per-frame segment labels, which are then used to assess the performance of the subject for the given task.

### 3.1 RGB Image Pre-processor

The transformer, the HMM, MSTCN++, and the rule-based decision tree pipelines have a feature extractor as demonstrated in Fig. 3 that converts the raw RGB frames into suitable input features. The frames capture the activity space that includes the patients upper body, the objects and the activity mat. All 12 tasks in the SARAH system involve movement in relation with one or two objects. Therefore, it is imperative to detect key movement features associated with both the patient’s upper body and the objects.

#### 3.1.1 Patient Skeleton Detection

We use OpenPose (Cao et al., 2019), an open source pose estimation technique, to extract 2D patient skeletons from videos. OpenPose generates 135 keypoints per-frame that include 25 body keypoints [4(A)], 21 keypoints for both hand [4(B)] and 70 keypoints for the face. These keypoints are the (*x, y*)-pixel coordinates of the skeleton joints as shown in Fig. 4(C). In the presence of multiple persons in the captured frame, we measure the area of the estimated upper body skeleton and consider the person with the largest area. We find that this simple pre-filtering approach worked in all cases within our test set to gives us the actual patient’s keypoints. In the SARAH system, the side camera used in this experiment always focuses on the impaired arm and the non-impaired arm is obscured. Therefore, we exclude the estimated keypoints from the non-impaired limb in our analysis framework. Since we are only interested in the upper body keypoints, we only consider keypoints above the lower torso line (keypoints 9, 8, and 12). Also, because of the non-standardized placement of the camera in this experiment, the face was not always within the frame and therefore, we excluded the face keypoints from further analysis. After all the exclusions, we use the two sets of keypoints as indicated in Fig. 4(A). The set of keypoints inside the red circle is for right-hand impaired patients and the blue one for the left-hand impaired patients. All keypoints are normalised with respect to the image frame.

**Figure 4.**
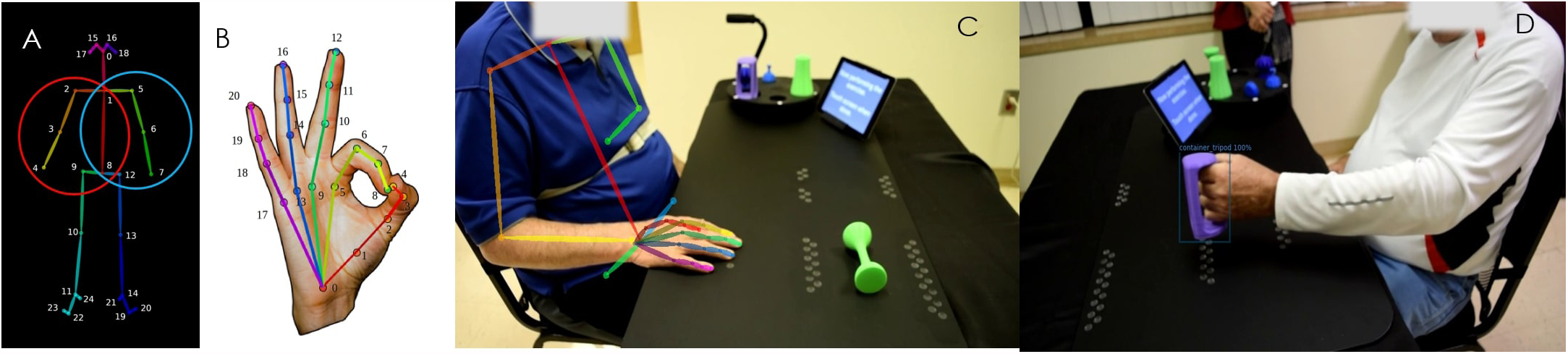
(A) 25 body keypoints that OpenPose generates following the COCO dataset; the circles indicate the set of keypoints used in the porposed analysis framework for the right hand imapired patient (red circle) and the left-hand impaired patient (blue circle) (B) 21 hand keypoints (C) OpenPose extracted upper body skeleton overlapped on the actual frame (D) Detected bounding box and object label using Faster RCNN

#### 3.1.2 Object Detection and Tracking

It was also imperative to obtain high quality object identification and location data for every frame, as the type of object being used and the relationship between the objects and the keypoints extracted from the subjects play an important role in determining segment classification. To this effect, we fine-tuned an object detection algorithm which classifies the objects in the frame, places a bounding box around the classified object, and provides the object/bounding box location relative to the overall frame. We considered a Faster-RCNN model (Ren et al., 2015) pre-trained on MS COCO (Lin et al., 2014) dataset for our experiments. To fine tune this object detection model we first labeled all objects being used across the 12 performed SARAH tasks from a small number of videos using CVAT (), an open-source annotation tool. In total we had a training set of around 23,253 training images and we validated it on 11,112 images with a mean IOU score of 0.67. We then used the open source detectron 2 framework (Wu et al., 2019), built on top of PyTorch (Paszke et al., 2019) to fine-tune the model.

As stated in Section 2.1, there is a high variability in the training data, owing to issues such as use of a single camera view, non-standardized camera angles, variability in the patient’s movement, unconstrained ways of grasping the objects, errors in transportation etc. As a consequence, the object detection model misses certain objects in certain frames due to factors such as occlusions or dropped objects, and sometimes even misclassifies certain objects due to partial visibility. However, as described in Section 1, the task being attempted in each captured video is known to the algorithm and each task is associated with specific objects. We use this knowledge to filter out misclassifications. To further improve the smoothness of the trajectories of the objects, we use a simple Kalman filter based tracking algorithm SORT (Bewley et al., 2016). Given the fine-tuned object detection model, the pipeline constructed is as follows. First, we extract all the frames in the video as images. Second, the trained Faster-RCNN is run on each of the frames thus collecting per-frame object data. Third, misclassifications are corrected automatically and the trajectories of the tracked objects are smoothed.

#### 3.1.3 Denoising Pose Keypoints and Object Locations

The activity space in different videos has high variability. As the camera angle and patient position are different for each captured session, the ratio of the activity space to the frame resolution changes. Low camera resolution and low frame-rate (used so as to allow efficient capture of long sessions), variable focus (cameras were set up by non-experts without standardized instructions) and varied lighting conditions (typical of therapy sessions happening in different contexts), create challenges for pose-estimation methods from the raw frames. As a result, the keypoint detection and bounding box estimation accuracy can suffer, sometimes even completely missing all keypoints and not detecting any bounding box in a frame. To reduce the missing data problem we first detect the outliers. We calculate the z-score for each sample using,

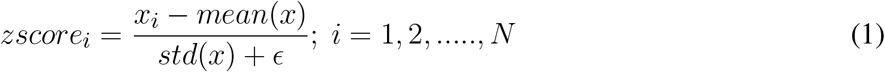

where *x*_*i*_ is the keypoints or object locations for the *i*^*th*^ frame (an *ϵ >* 0 is used to guard against pathological situations). If the z-score value is higher than a threshold, we consider the sample as an outlier. For our case, using a threshold of 2.5 gives good performance. Then we use spline interpolation to fill the missing values or replace the outliers.

#### 3.1.4 Normalization of input data

Denoising and normalizing the input data is a vital step in achieving good performance through machine learning. We experimented with three different techniques to find the best techniques for normalizing both the patient keypoints data and the object locations-(i) global normalization with respect to the image frame (ii) normalization with respect to the activity mat area and (iii) normalization using a computed homography matrix.

##### Global-Norm

Here, we just divide the raw patient keypoints and object location data with the image frame’s width and height, thus mapping the data to a [0, 1] range. In practice due to the high amount of noise in the data, and variability in inter-patient performance, this global normalization fails to achieve good results and was therefore not applied (see Table 6).

##### Mat-Norm

Here, we normalize the data with respect to the ratio of the area of the activity mat to the total image area. This is done by labeling the activity mat in one of the frames per subject per capture session. For every capture session the camera position is set at the beginning. We could thus precompute the area of the mat for all recorded tasks of each capture session. This normalization with regards to the mat area removes a good degree of variability and thus performs better than global normalization.

##### Homography-Norm

Since the camera angles across subjects and across attempts vary considerably, it is important to reduce the impact of this variation. To achieve this, we apply a simple homography-based re-projection of the object and keypoint data. First, a homography matrix is pre-computed that transforms data from image frame to the coordinate frame defined by the activity mat. This transformation matrix is then applied to the object and keypoint data. Even though this achieves high invariance, it also removes discriminative features and thus there is not any observably significant improvement in the performance of the transformer models.

### 3.2 HMM features and algorithmic pipeline

#### 3.2.1 Kinematic Feature Extractor

Using the denoised keypoints and object locations, we calculate 20 kinematic features that, based on our prior work (Chen et al., 2011b), can be combined to successfully assess functionality and movement quality in upper extremity stroke rehabilitation. These features can be used to analyze the movement of the object, the affected limb and torso, and the patterns of human-object interaction. These 20 features are summarized in Table 1. The variability in stroke survivor movement and the low quality capture resulted in 7 of the 20 features being too noisy to support automated analysis. We therefore only used the 13 less noisy features for further analysis in the paper. In addition to these 13 features, we also calculate derivatives of some of the features. The derivatives introduce oscillations unique to segments and by learning those patterns the efficacy of the HMM in per-frame segmentation improves. A detailed description of the kinematic features are given in the supplementary section.

**Table 1.** List of kinematic features extracted from patient 2D keypoints and object co-ordinates Patient Features Object Features

| Patient Features | Object Features |
| --- | --- |
| limb Location | Object Location |
| limb Velocity | Object Velocity |
| limb Absolute velocity | Object Absolute velocity |
| Distance between limb and Torso Line | Distance between two Objects |
| Pronation and Supination | Object Angular Velocity |
| Relative angle between torso line and limb | Distance between Object and limb |
| Hand Openness | Distance between Object and torso line |
| Torso sway, rotation and lean |  |
| Elbow Extension |  |
| Jerkiness |  |
| Absolute limb angle |  |

#### 3.2.2 Hidden Markov Model with Prior Transitions

A hidden Markov model (HMM) (Rabiner, 1989) consists of a set of states, a transition model, and an observation model. Denote *S* = *S*_1_, *S*_2_, …, *S*_*N*_ as the *N* states. Evolution between states is modeled by the transition probability conditioned on the previous state. The transition probability between states is represented by a matrix, *A* = *a*_*ij*_; where *a*_*ij*_ = *Pr*(*S*_*j*_(*t* + 1)|*S*_*j*_(*t*)). For a given observation, *O* = *O*_1_, *O*_2_, …, *O*_*k*_, the emission probability distribution is represented by a matrix *B* = *b*_*j*_(*k*); where *b*_*j*_(*k*) is the probability of generating observation *O*_*k*_ when the current state is *S*_*j*_, where *k* is the total number of observation symbols.

HMMs have been used in movement quality assessment because of their good performance in detecting subtle inconsistencies in the movement (Osgouei et al., 2018; Nguyen et al., 2016; Deters and Rybarczyk, 2018). However, the topological structure of the HMM in most cases cannot be automatically determined due to the highly variant and small dataset (Deters and Rybarczyk, 2018). We therefore use expert knowledge based design constraints to model the topological structure of the HMM for different exercises. We have modeled 5 different transition matrices demonstrated in Fig. 5 that are based on the number of unique segments present in the 12 SARAH tasks. These matrices provide a strong prior as they are modeled after the therapist’s approach to parsing movement and focusing attention on key composite movement features per type of segment.

**Figure 5.**
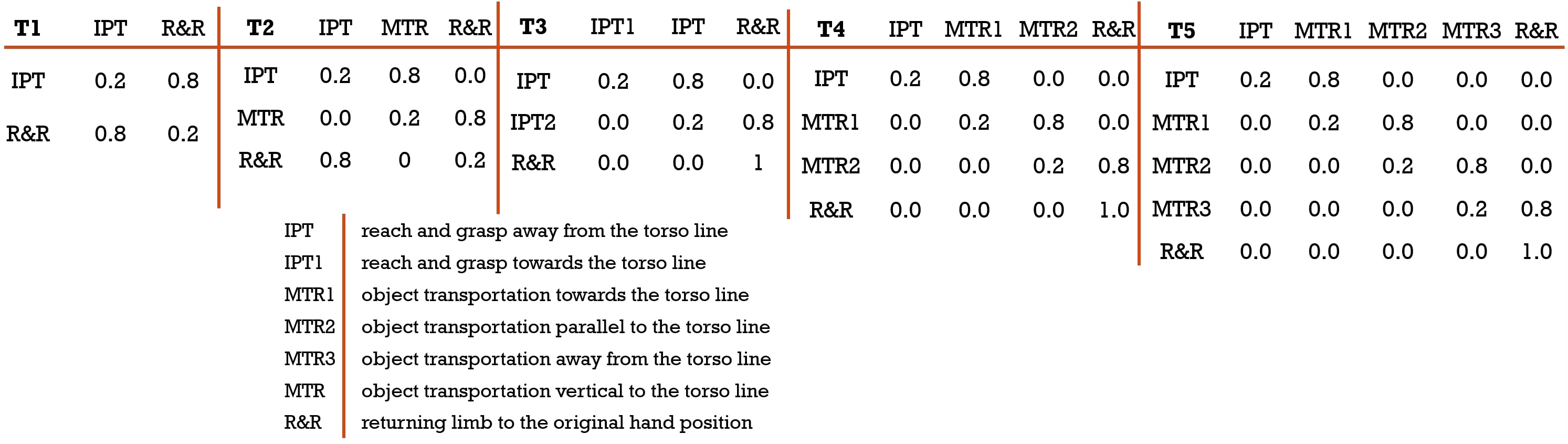
Different transitions matrix based on expert knowledge; (from left) T1: a 2× 2 transition matrix designed for task 1 and 2; T2: a 3 × 3 transition matrix designed for task 6,7, and 10; T3: a 3 × 3 transition matrix designed for task 9; T4: a 4 × 4 transition matrix designed for task 3,4, and 5; T5: a 5 × 5 transition matrix designed for task 8

**Figure 6.**
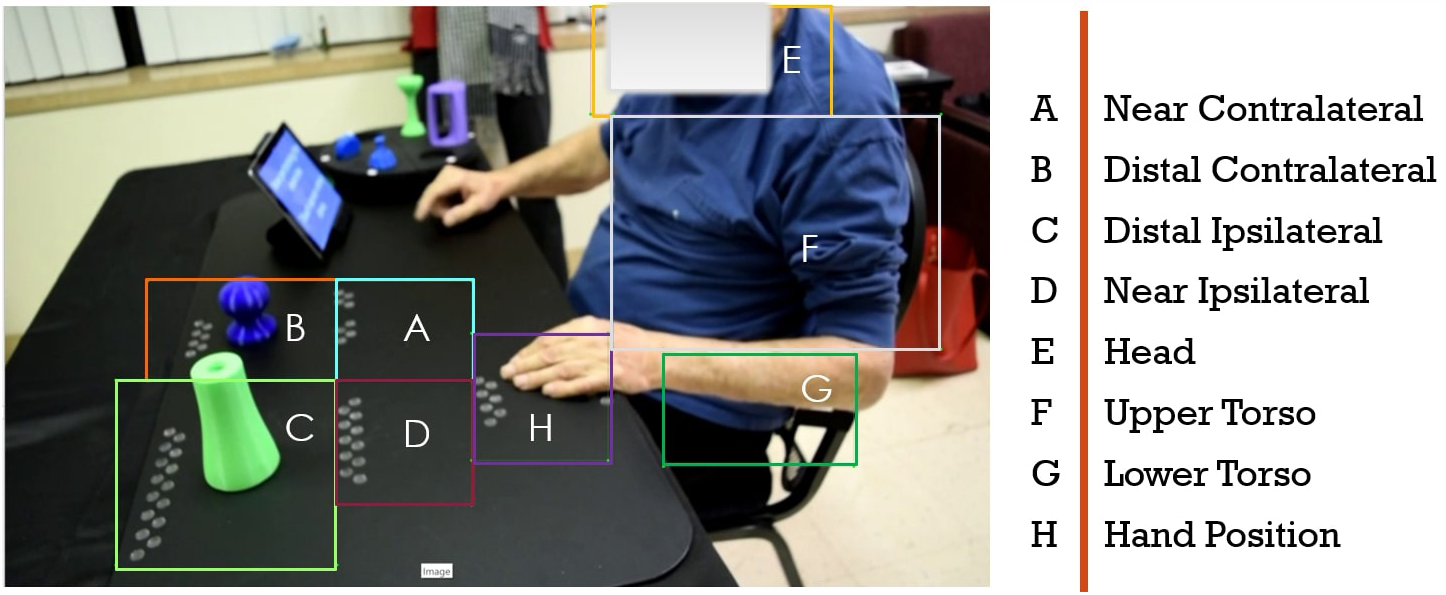
Drawn bounding boxes on the activity space and patients upper body; there are five bounding boxes on the mat and three on the patients body

To train the HMM, we first need to define the initialization for each state in the transition matrix. For example, to initialize an HMM with T4 transition matrix, we need 4 states: *S* = {*IPT, MTR*1, *MTR*2, *R*&*R*} . Each state is initialized with a normal distribution of mean, *µ* = 0.5 and standard deviation, *σ* = 0.1. We choose the normal distribution because it produced the lowest chi-square error after intensive experimentation with different distributions like beta, exponential, gamma, log-normal, normal, Pearson3, triangular, uniform, and Weibull. The distribution parameters can be randomly initialized, and we find that the model is not sensitive to random initialization. The training of the HMM tunes the parameters which are then compactly represented as,

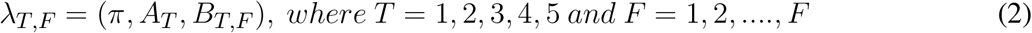

Here, the subscript *T* and *F* refer to different transition matrices and features and *π* is the initial state distribution and is defined as *π* = *π*_*i*_ where *π*_*i*_ is the probability of state *S*_*i*_ being the initial state. Based on the tasks, *π* = [1, 0, …]. Since all the tasks start with the state IPT, the first state probability is always 1 followed by zeros for other states present in the transition matrix. This is another strong design prior we leverage, in the form of supervision of the labels with state names. In this case, maximum likelihood estimation (MLE) estimates the emissions from data partitioned by the labels and the transition matrix is calculated directly from the adjacency of labels. Then, various transition matrices and their initialization are further tuned with patient data. As we provide more training data, the final models are expected to become more generalizable across different types of tasks. Another prior we use for structuring our HMM relates to the use of distributions of individual kinematic features. Therapists establish through experience an expected distribution per feature per segment (i.e. Gaussian velocity profile for IPT).Therefore, to learn the distribution parameters for each of the kinematic features, it is a requisite to train separate HMMs for each of those features.

We use the open source toolbox Pomegranate (Schreiber, 2018) for the training and estimation of the state probabilities for each feature. Then, we formulate an objective function on the state probabilities of the kinematic features to estimate one set of state probabilities. For this purpose we use a mean squared error based objective function defined as,

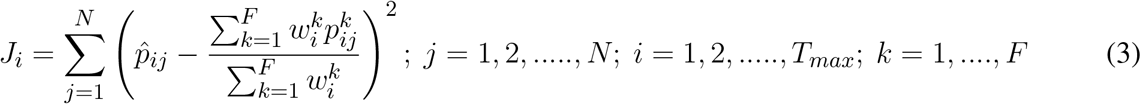

where, 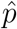 denotes the per-frame categorical labels and *p* is the predicted state probabilities. *N* is the total number of observations in the training set, *T*_*max*_ is the maximum observable frame number, *F* represents the total number of features, and *w* is the weight for optimization. We optimize the objective function by adding two constraints on the weights. Firstly, restricting the range of weights *w* ∈ [0, 1] and secondly, constraining the weights to unit sum: 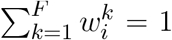. The objective function was minimized using Sequential Least Square Programming (SLSQP) (Nocedal and Wright, 2006) due to its ease of implementation and because our objective function naturally is an instance of this type of problem.

### 3.3 Transformer Pipeline

In addition to HMMs, we also explore deep-learning based transformers Vaswani et al. (2017), which have attracted a large amount of interest in the natural language processing community due to their strengthes in modelling long term dependencies while being computationally efficient and avoiding problems such as vanishing gradients in other deep-learning based time-series approaches like long short-term memory (LSTM) based approaches. Since our goal is video-segmentation, and per-frame classification, we model this problem as a Sequence-to-Sequence problem (Sutskever et al., 2014), where the input is a multivariate time-series determined by body keypoints and object data. Given this data, we output discrete segment labels for every timestep (frame). The pipeline is as follows: (i) concatenate normalized pre-processed zero-padded keypoints and object location data to obtain 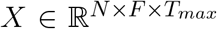 where *N* is the number of videos, *F* is the total number of feature and *T*_*max*_ is the maximum observable frame number including the zero-padding, (ii) partition the data *X* into training *X*^*tr*^, validation *X*^*val*^, and test *X*^*te*^ sets randomly, with a constraint that no patient-task pair should be repeated in train and test sets, (iii) train the transformer with the objective to find a function *g* that maps *X*^*tr*^ to *Y*, where *Y* ^*tr*^ ∈ ℝ^1*×T s*^ is the per-frame label vector, by minimizing cross entropy loss between *f* (*g*^*tr*^) and *Y* ^*tr*^. We train the network with different augmentations on the input data such as random drift, Gaussian noise etc using (Arundo, 2019) to avoid overfitting.

#### 3.3.1 Transformer Architecture

We adopt the transformer architecture from (Vaswani et al., 2017), with minor modifications to adapt it for timeseries (Cohen et al., 2020): (i) we replace the embedding layer with a generic layer; and (ii), we apply a window on the attention map to focus on short-term trends because the segments are short. Other hyperparameters are as follows: number of encoder-decoder pairs are set to 2, number of heads as 4, the dimension of key query and value and an attention window of 100 along with sinusoidal positional encoding.

### 3.4 MSTCN++ Pipeline

Temporal convolutional networks (TCNs) (Lea et al., 2017) are shown to outperform methods such as LSTMs on various time-series modeling problems. Building on the success of TCNs, Farha et al recently developed MS-TCN (Farha and Gall, 2019) a multi-stage variant, by stacking multiple TCNs. This multi-stage approach is shown to outperform TCNs for video segmentation. We have thus used MS-TCN (Farha and Gall, 2019) and its variants MS-TCN++ (Li et al., 2020) for our experiments. While our transformer and HMM based methods operate on carefully constructed features from keypoints and objects, our TCN approach can directly use features extracted by a feature-learner approach such as a pre-trained video classification network, I3D (Carreira and Zisserman, 2017). I3D is trained on the Kinetics dataset (Kay et al., 2017), which contains videos of human actions, with classes such as single person actions, person-person actions, and person-object actions. Since the I3D model that has been trained on this large scale dataset, features extracted though the model can capture representations of complex human activity, including human-object interaction, thus making the model a fitted choice for composite feature extraction for our analysis. We train MS-TCN++ on the features from the I3D model on our data with an objective to minimize the weighted combination of (i) cross entropy loss between the predicted segment classes and ground truth, and (ii), a penalty for over segmentation. We fix the multiplier to the penalty term, which is a hyperparameter at 0.15 as mentioned in (Li et al., 2020). We train the network for 300 epochs with an ADAM optimizer (Kingma and Ba, 2015), learning rate of 10^*−*3^ and step-wise reduction in learning rate by a factor of 0.8 for every 30 epochs.

### 3.5 Weighted Average Ensemble

Finally, we fuse the outputs of the data driven models with the expert knowledge constrained models. The HMM algorithm uses kinematic features and design priors that are constrained by the expert knowledge. The predictions from the HMM encode information about the segments and task level of the movement analysis hierarchy shown in Fig. 1. On the other hand, techniques like Transformer and MSTCN++ use kinematic and composite features to perform better in the frame level but fail to encode information about the segments and the tasks. To combine the strengths of each approach, we fuse the analysis outcomes of the different algorithms at the level of the segment. As discussed in section 2.1, the segment level of the hierarchy is expected to have the highest level of agreement between expert movement analysis and computational analysis. We implement this hypothesis by taking a weighted average ensemble of the state probabilities from each model. If 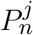 is the probability of *j*^*th*^ state of model, *n*, then the predicted state is calculated as,

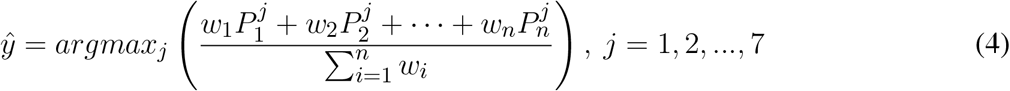

here, *w*_*i*_ is the weight for model *i* and *ŷ* is predicted states. We have conducted a grid search for each weight in the range of [0.1, 0.2, 0.3, …, 1.0]. Thus for a three model ensemble, we run the evaluation 10 × 10 × 10 = 1000 times for each combination of weights. We experimented with 4 ensemble schemes to find the combination that provides the most robust results: (i) HMM and Transformer, (ii) HMM and MSTCN++ (iii) Transformer and MSTCN++ and (iv)HMM, Transformer, and MSTCN++.

### 3.6 Block Based Task Completion Assessment

To perform task completion assessment we need to organize the per-frame segment labels into segment blocks. We again leverage therapist expertise for grouping segment samples into segment blocks (as embedded in the SARAH system design), to automatically extract segment blocks and assess the order, duration, and continuity of the blocks. To calculate segment blocks, we first denoise the ensemble predictions. The per-frame segment predictions resulting from the ensemble model have two types of noise: false transitions and missing transitions.

#### 3.6.1 Denoising of Ensemble Predictions

In the SARAH system, the performance of all tasks is realized through a sequence of a small number of segments. These segments are performed with a defined set of objects, and within a defined activity space. Further, they are executed at a speed that is constrained by human biomechanics and potentially constrained by stroke induced impairment. We can filter some of the false predictions through a Gaussian filter for each transition model that implements these priors. In Fig. 10, we illustrate the per-segment filters for a 3 × 3 transition matrix model. The filters can be mathematically expressed as *G*_*s*_(*µ*_*s*_, *σ*_*s*_), where *µ*_*s*_ and *σ*_*s*_ denotes to the mean duration and standard deviation of each segment, *s* calculated from the training data. For a test data of length, *N*, we calculate the posterior distribution and use a 75% confidence cutoff to get the upper and lower bounds for each segment. We only accept a predicted transition, if it falls within the bounds. In addition, we pose a 20 frame window constraint on the predictions since all the task segments in the SARAH system require more than 1 second to complete. Therefore, any transitions within 20 frames is discarded. This filtering technique reduces the majority of the false predictions.

#### 3.6.2 Rule-Based Binary Decision Tree (RBBDT)

To correct the remaining false predictions and find the missing segment transitions, we use a rule-based binary decision tree (RBBDT) to encode the process that the experts (therapists) use to segment the patient movement. As observed during the development of the SARAH system with therapists, instead of classifying each frame of movement, therapists utilize a few key events to find transition point candidates between segments. These events are primarily based on three relationships: (i) patient -object interaction, (ii) patient -activity space relationship, and (iii) object -activity space relation. People execute functional tasks differently (i.e. the trajectory of raising a glass to our mouths to drink is slightly different for each person). These differences are much more pronounced for stroke survivors as they have different types of impairment. Therefore, therapists organize the patient activity space into a few generalizable regions that are robust to variation. Standardized stroke rehabilitation assessment tests (i.e. WMFT, ARAT) rely on such generalizable regions and the regions have accordingly been adopted in the design of the SARAH system (see section 1).

To model relationships between the object(s), patient, and the generalizable activity regions, we recreate the activity regions as eight bounding boxes on each frame of the video. These bounding boxes are illustrated in Fig. 6. Our approach is similar to region based object detection (Ren et al., 2015) techniques, where the bounding boxes are created based on the relationships between different objects and the space. Our bounding boxes, divide the body into the regions of head, upper and lower torso, since movements of the impaired arm towards the head (i.e. for feeding) have different functionality than movement of the impaired limb towards the torso (i.e. dressing). Moving the end point of the impaired upper limb towards the upper and lower torso presents different challenges for different patients. The tabletop activity space is divided into an ipsilateral and contralateral area since the engagement of these two spaces requires different coordination of joints and muscles. The ipsilateral and contralateral areas are further divided into proximal and distal areas, since different extension patterns of the arm are needed to engage the distal space (see Fig. 6). We use these eight bounding boxes to calculate the space-patient, space-object, and patient-object relationships that appear in Table 2.

**Table 2.** List of RBBDT features calculated using patient keypoints, object co-ordinates and center of 8 bounding boxes

| Index | Feature notation | Definition |
| --- | --- | --- |
| 1 | $d_{Fl}$ | distance between upper torso, F and the limb |
| 2 | $d_{Gl}$ | distance between lower torso, G and the limb |
| 3 | $d_{El}$ | distance between head, E and the limb |
| 4 | $d_{hl}$ | Distance between limb and hand bounding box, h |
| 5 | $d_{Al}, d_{Bl}, d_{Cl}, d_{Dl}$ | distance between limb and bounding box A,B,C, and D |
| 6 | $d_{Fo}$ | distance between upper torso, F and the object |
| 7 | $d_{Go}$ | distance between lower torso, G and the object |
| 8 | $d_{Eo}$ | distance between head, E and the object |
| 9 | $d_{ho}$ | Distance between object and hand bounding box, h |
| 10 | $d_{Ao}, d_{Bo}, d_{Co}, d_{Do}$ | distance between object and bounding box A,B,C, and D |
| 11 | $d_{lo}$ | distance between limb and object |
| 12 | $d_{oo}$ | distance between two objects |
| 13 | $\frac{d}{dt}()$ | first derivative of the above features |

We use previously calculated object and end point position and velocity to mark the object and limb end point stopping time. We can then calculate multiple distance features like limb-bounding box distance, object-bounding box distance, and limb-object distance. These simple distance features do not exhibit a simple statistical pattern (e.g. Gaussian) across different patients for each type of segment and are therefore not easily amenable to training the HMM segmentation algorithm. However, under the RBBDT framing they did indeed become usable for detecting segments blocks because they encode the features used by experts to define segment blocks as shown in Table 2.

In our previous work (Venkataraman et al., 2014), we used a standard classification and regression tree (CART) for modeling therapist decision processes in movement assessment. Data driven tuning of the parameters of these decision trees is sensitive to noisy data. Since the dataset for this experiment is small and noisy, we developed an approach for manually tuning the thresholds and split branches, and selecting the order of features of the RBBDT. Our manually tuning is supported by training data. The goal of the tuning is to place the features in descending order of observability and error for each type of transition prediction with the most observable and accurate feature coming first in the decision sequence. The exercises of the SARAH system have different combinations of segments and variable object locations. Therefore, the RBBDT based prediction of each segment transition, the transition between two segment blocks, requires a unique order of features. In the appendix section, we show the order of features used for each transition of the different SARAH tasks based on the mathematical notation from 2. If a candidate sample of a feature stream meets the threshold condition, the confidence value of that candidate as a transition point increases.

Let’s consider the previous example of exercise three and four where the patient transports the object close to their mouth to simulate drinking action. To get the candidates for the IPT-MTR1 transition, the highest observable feature is the location of the object and limb as a phase change occurs at the beginning of MTR1. The next most observable feature is the directionality of the object’s movement in relation to the bounding boxes. The process for calculating the confidence value of the candidates for the above IPT-MTR1 transition is illustrated in Table 3. In the shown example, RBBDT generates a possible candidate when two or more conditions are true for the sample. Therefore, we set the confidence threshold to 0.5. It is important to note, that the RBBDT does not aim for accuracy at the per-frame classification level as this accuracy is secured through the ensemble model presented above. The RBBDT aims to only find transition points organizing the data stream into segments blocks that are feasible (could have been performed by a stroke patient), while minimizing missed transitions and false transitions between blocks.

**Table 3.** Confidence calculations based on binary decision from the rules

| Object phase | limb phase | bounding box Directionality | Confidence, C |
| --- | --- | --- | --- |
| 0 | 0 | 0 | 0 |
| 0 | 0 | 1 | 0.25 |
| 0 | 1 | 0 | 0.25 |
| 1 | 0 | 0 | 0.25 |
| 1 | 0 | 1 | 0.5 |
| 1 | 1 | 0 | 0.5 |
| 0 | 1 | 1 | 0.5 |
| 1 | 1 | 1 | 0.75 |

#### 3.6.3 Task Completion Assessment

For a given task, the segment blocks will inform us about the continuity and order of the segments. In addition, we can calculate the duration of the completed task. Once we have calculated the segment blocks and duration for each task, we can compare that calculation to the expected sequence of segments and duration for each task for a coarse assessment of task completion. The algorithm checks for three conditions:

- Is the task completed with all the necessary segment blocks?
- Is the order of the segment blocks correct?
- Is the task performed within allowable time?

If a task satisfies all three conditions, the algorithm gives a score of 3. If the first two are satisfied then it generates a score of 2. If either of the first two are not satisfied then it generates a score of 1. Finally, a score of 0 is assigned if the task is not attempted. Although these scores are similar to the scores assigned by therapists they do not take into account issues of movement quality. Therapists may assign a score of 2 to a task that is completed within the allowable time because of issues of movement quality. Since our methodology connects different types of movement features to assessment of segment and task execution, in the future we plan to use actual therapists ratings of patient videos to further train our algorithms so they can automatically assess task completion and movement quality.

## 4 EXPERIMENTAL SETTINGS

Our initial dataset consists of 610 captured videos. The experimental dataset includes 404 videos after exclusion of videos where severe limb impairment caused multiple object dropping or multiple segment occurrence leading to an incomplete exercise. Including these videos would severely skew the training sets at this early stage. However, some of these videos were included in the test sets for task completion and in the future these videos can be included in the training sets too. In our dataset each patient performed each task multiple times in a single session. Therefore, we design random split experiments based on three factors: patient ID, session ID and task number. For any task number, the same patient ID with the same session ID is included in either of the training or test set but not in both. We choose different random seeds and create 5 random experiments for fairness based on this selection method. The average training and test size of the 5 splits are 370 and 34, respectively. We implement the deep learning experiments on two NVIDIA RTX 2080 Ti GPUs. To evaluate the segmentation performance of the experimented models, we calculate frame-wise accuracy, precision, and recall. The calculation formula for the matrices are given in the appendix section. For each of the split experiments, we evaluate the matrices independently and calculate mean and standard deviation of the results for all the algorithms.

## 5 RESULTS

### 5.1 HMM Segmentation Results

Table 4, shows the segmentation results using the proposed five-transition-matrix HMM. The average per-frame accuracy is 77.82%±2.88 meaning around 78% of the frames are labeled correctly. The precision and recall values are 78.60%±2.47 and 78.26%±2.76, respectively. Additionally, we have performed the following ablation studies to demonstrate the efficacy of the proposed five-transition-matrix HMM.

**Table 4.** Per frame segmentation Results using the Hidden Markov Model

| Split Index | Frame Wise Accuracy | Precision | Recall |
| --- | --- | --- | --- |
| 1 | 75.52 | 77.94 | 76.04 |
| 2 | 78.95 | 79.21 | 79.08 |
| 3 | 80.75 | 80.17 | 80.96 |
| 4 | 74.08 | 74.69 | 74.72 |
| 5 | 79.82 | 81.01 | 80.49 |
| Mean | 77.82 | 78.60 | 78.26 |
| STD | 2.88 | 2.47 | 2.76 |

#### 5.1.1 Number of features

We experimented with the proposed HMM using various numbers of features. The feature selection is an important factor as the combination of correct features captures the unique patterns of different transitions. To understand the effect of feature selection, we perform three experiments: (i) 13 kinematic features; (ii) six kinematic features including four derivatives; and (iii) one composite feature. To get the composite feature from 13 kinematic features, we experimented with different dimensionality reduction techniques like PCA, NMF, LDA, and RP (Abdi and Williams, 2010; Choi, 2008; Balakrishnama and Ganapathiraju, 1998; Bingham and Mannila, 2001). Based on the performance, we choose PCA to reduce the dimension of 13 features into one and produce the composite feature. In Fig. 7 (a), we illustrate the comparative results for the above three experiments. As evident from the illustration, the best performing result was achieved when a combination of the raw kinematics and the derivatives were used as input to the HMM.

**Figure 7.**
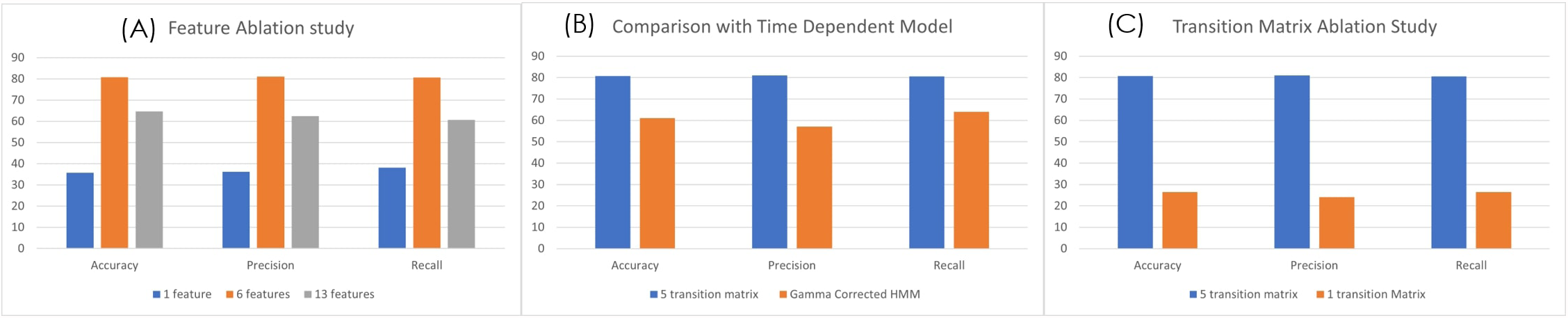
(a) Feature ablation study using 13 and 6 kinematic features and 1 composite feature (b) Comparison between the 5 transition matrix vs 1 transition matrix model HMM (c) Comparison between 5 transition matrix HMM with 5 transition matrix time dependent HMM model

#### 5.1.2 Transition matrix

The proposed HMM model uses five transition matrices. We also experimented with one transition matrix model to understand the capability of the HMM to learn different variations of segment transitions across all types of tasks. A 6 × 6 transition matrix is necessary to represent all the segment transitions across all performed tasks in our data set. In Fig. 7 (c), we show the comparison between the one transition matrix HMM and the proposed five transition matrix HMM (with each transition matrix calibrated to particular types of tasks).

#### 5.1.3 Time Dependency

We also experimented with time dependent models and wanted to exploit the duration of segments to correct the predictions from the HMM. To model time dependency in the HMM predictions, we use the gamma correction (Turaga et al., 2009) on the predictions of the HMM. We calculate the gamma probabilities using:

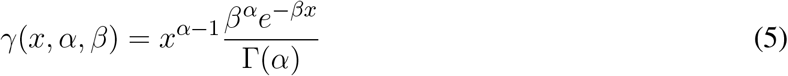

where, *α* and *β* is calculated per segment using mean (*µ*) and standard deviation (*σ*) of the segments in the training set. Since the dataset has a very high standard deviation, the gamma function (Γ(*α*)) results in a large value and thus Eq. (5) generates probabilities *<* 0.01. Therefore, the effect on the HMM predictions are negligible. In Fig. 7 (B), we illustrate the comparison with the proposed 5 transition matrix HMM model.

### 5.2 Transformer Segmentation Results

Like the HMM, the transformer was also tested on five random splits. In Table 5, we demonstrate the segmentation results for the 5 splits. As described in section 3.1.2, we experimented with two types of normalization technique to remove activity space variance. We present the result for both normalization techniques in Table 5. We conducted two additional studies to understand the dynamics of our transformer pipeline by ablating on (i) the number of features used as input; and (ii) the amount of labeled training data needed. The former ablation study attempts to identify the features that are beneficial, while the latter answers the reduction in time and cost needed to label the data by hand.

**Table 5.** Per frame segmentation Results using Transformer with two different type of activity space normalization

| Normalization | metrics | 1 | 2 | 3 | 4 | 5 | Mean | STD |
| --- | --- | --- | --- | --- | --- | --- | --- | --- |
| Homography-Norm | ACC | 70.07 | 74.29 | 75.85 | 65.34 | 72.37 | 71.59 | 3.67 |
|  | Precision | 65.03 | 61.68 | 68.69 | 58.37 | 63.20 | 63.40 | 3.43 |
|  | Recall | 61.84 | 61.45 | 68.08 | 55.55 | 60.54 | 61.49 | 3.99 |
| Global-Norm | ACC | 69.60 | 78.55 | 75.29 | 75.36 | 76.20 | 75.00 | 3.29 |
|  | Precision | 68.62 | 69.43 | 72.58 | 72.40 | 70.83 | **70.77** | 1.76 |
|  | Recall | 65.37 | 68.88 | 72.77 | 69.51 | 68.73 | **69.05** | 2.63 |
| Mat-Norm | ACC | 73.43 | 78.82 | 78.55 | 73.67 | 77.24 | **76.34** | 2.34 |
|  | Precision | 70.20 | 72.35 | 73.49 | 66.72 | 67.74 | 70.10 | 2.59 |
|  | Recall | 68.71 | 70.55 | 74.19 | 61.94 | 66.43 | 68.36 | 4.10 |

**Table 6.** Ablation results on Transformers using different number of keypoints. Using only wrist keypoint outperforms the rest.

| # keypoints | ACC |  | Precision |  | Recall |  |
| --- | --- | --- | --- | --- | --- | --- |
|  | Mean | STD | Mean | STD | Mean | STD |
| **1** | **76.34** | 2.34 | **70.10** | 2.59 | **68.36** | 4.10 |
| 2 | 69.65 | 1.66 | 55.34 | 3.67 | 53.87 | 4.02 |
| 4 | 71.30 | 3.20 | 67.51 | 4.23 | 66.14 | 2.89 |
| 8 | 67.80 | 2.53 | 60.19 | 7.37 | 56.47 | 6.15 |

#### 5.2.1 Number of Features

We experimented with different number of keypoints while always using the object features as input to the transformer based pipeline. We decreased the number of keypoints from 8, 4,2 and then to 1. We selected the keypoints for each reduction based on our prior work (Baran et al., 2015) that defined the relative importance of different key points in characterizing upper limb impaired movement. The best performing result was achieved using just the wrist keypoints and object locations. This conforms to the observations made in (Baran et al., 2015). The results are given in Table 6.

#### 5.2.2 Amount of Training data

. We experimented with reducing the amount of training data to the transformers to better study the generalization properties of our pipeline in low data scenarios. This is a practical setting, as the amount of time required to label the data is very high. However, as transformers contain large number of parameters, it is important to study the over fitting trends as well. We report average accuracies on the same test data for fair comparison while changing the size of input training data. As can be seen from the Table 7, if the amount of data available is too small, unsurprisingly the performance is very poor due to overfitting; however, as size of available data increase, the performance improves significantly plateauing after sometime. It is encouraging that even with 25% reduction in training data(404 to 300) the drop in performance is only 2% thus showing the robustness of the model.

**Table 7.** Transformer performance with varying sizes of training data. As expected, more data prevents overfitting thus explaining the better performance.

| # of samples | ACC |  | Precision |  | Recall |  |
| --- | --- | --- | --- | --- | --- | --- |
|  | Mean | STD | Mean | STD | Mean | STD |
| 100 | 65.00 | 1.85 | 57.23 | 3.12 | 55.12 | 3.05 |
| 200 | 71.97 | 3.00 | 64.63 | 1.79 | 62.85 | 2.17 |
| 300 | 75.54 | 2.57 | 69.49 | 2.54 | 68.11 | 3.88 |
| 404(all data) | **76.34** | 2.34 | **70.10** | 2.59 | **68.36** | 4.10 |

### 5.3 MSTCN++ Segmentation Results

The results for the five split experiment using the MS-TCN++ is shown in Table 8. This is the best standalone model in the pipeline as evident from the values. All coordinates were normalized with respect to the activity mat area as described in 3.1.3 refered to as Mat-Norm. As can be seen from 8, MS-TCN++ achieves the best performance in terms of per-frame classification error. However, as will be shown in section 5.5, even though it achieves better classification error, it performs poorly in standalone estimation of the completion of the task. As MS-TCN++ is a completely data driven model it fails to encode any design priors or expert knowledge (i.e. expected order of segments). Thus, the few errors that MS-TCN++ makes are highly harmful. As an example, while every task should start with an ‘IPT’ segment, the MS-TCN++ based model labels the first few frames of some tasks as ‘MTR’ thus affecting the automated task completion performance. Thus, for higher level automated decisions (assessing completion of segments and tasks by patients) it is imperative to fuse the MS-TCN++ with other models that encode prior knowledge such as our implementations of the HMM and transformer based models.

**Table 8.** Per frame Segmentation Result using MSTCN++. This method outperforms HMM and transformers for the task of per-frame segmentation.

| split | 1 | 2 | 3 | 4 | 5 | Mean | STD |
| --- | --- | --- | --- | --- | --- | --- | --- |
| ACC | 79.75 | 85.96 | 83.26 | 79.96 | 80.60 | 81.91 | 2.38 |
| Precision | 76.84 | 85.29 | 82.05 | 74.30 | 75.08 | 78.71 | 4.25 |
| Recall | 77.64 | 84.04 | 82.24 | 74.68 | 75.51 | 78.82 | 3.70 |

### 5.4 Ensemble Model Segmentation Results

We experiment with three combinations of model ensembles. In Table 9, we illustrate the results of three ensemble models. The best performance is achieved when all three models are combined and that finding is consistent through all the splits. This is rather unsurprising because, the models are all complementary in nature with MS-TCN++ being completely data driven, while our transformer and HMM based models encode certain amounts of prior knowledge.

**Table 9.** Results using different ensembles. (i) Ensemble1: Transformer and HMM, (ii) Ensemble2: HMM and MS-TCN++ and (iii) Ensemble3: Transformer, HMM and MS-TCN++. We observe that ensemble3 outperforms the rest.

|  | Ensemble1 |  | Ensemble2 |  | Ensemble3 |  |
| --- | --- | --- | --- | --- | --- | --- |
|  | Mean | STD | Mean | STD | Mean | STD |
| ACC | 81.01 | 1.99 | 83.76 | 2.77 | 85.08 | 2.14 |
| Precision | 77.28 | 4.38 | 81.40 | 3.92 | 84.34 | 2.61 |
| Recall | 77.88 | 4.46 | 81.57 | 3.66 | 84.6 | 2.68 |

### 5.5 Task Performance Assessment Scores

In Table 10, we demonstrate the results of the block based segmentation. As evident from the table, with the incorporation of the proposed denoising and RBBDT, around 99% of the segment blocks can be labeled correctly. We also calculate task completion accuracy based on the continuity, and order of the segment blocks and duration of the tasks. A task is completed if it has an appropriate order of segments. As shown in Fig. 9, with the highest as 100% for split 4 and lowest as 87.5% for split 3, the average task completion accuracy is 92%. This means 92% of the tasks in the test case that were tagged as completed by the therapists are also labeled as “completed” by our algorithm. These are the tasks that have a score of either a 3 or 2. The algorithm generates a score of 2 only when a patient takes longer time to complete the task correctly. In Fig. 9, we show the percentage of correctly predicted 2s using the segment blocks for three experiments. For these experiments we varied the range of standard deviation (std). As seen from the figure, our algorithm predicted maximum 22% of the 2s among the videos that are rated 2s by the therapist. All these tasks are labeled as completed both by the algorithm and the therapist.

**Table 10.** Segmentation block results.

| split | Segment wise Accuracy | Precision | Recall |
| --- | --- | --- | --- |
| 1 | 99.33 | 99 | 98.5 |
| 2 | 98.48 | 98.18 | 97.04 |
| 3 | 98.66 | 98 | 97 |
| 4 | 100 | 100 | 100 |
| 5 | 98.66 | 98.5 | 97.5 |
| Mean | 99.03 | 98.73 | 98.01 |
| STD | 0.56 | 0.71 | 1.13 |

## 6 DISCUSSION

In this paper, we propose a hierarchical model for automatically segmenting stroke survivor’s movements and generating task performance assessment scores during rehabilitation training. The hierarchical model fuses expert knowledge-biased approaches with data-driven techniques. The expert knowledge is more observable in the higher layers of the hierarchy (task and segment) and therefore more accessible to algorithms incorporating high level constraints relating to activity structure (i.e. type and order of segments per task). The lower layers need to be addressed primarily through data driven techniques. By developing a sequence for combining complimentary algorithms that effectively encode the information from the different layers, we produce robust segmentation and task assessment results driven by noisy, variable, and limited data.

The MSTCN++ (Li et al., 2020) is a data driven technique that relies primarily on the RGB data layer of our hierarchy and on composite kinetic features (the features extracted through the pre-processor) that are trained through generic videos of human activity. Therefore, the composite features are not fully observable and are not directly related to upper extremity functional tasks. Furthermore, the algorithm cannot incorporate the higher layer constraints resulting from the expert driven design of the system (segment vocabulary, composite features related to each segment, order of segments per task). As the algorithm is most sensitive to the lower layers of the hierarchy, it performs the best in terms of per-frame segment classification, but because it does not incorporate higher layer constraints, is prone to errors in the order of segments. For example it can classify the beginning frames of a task as belonging to an MTR segment when all the SARAH tasks start with IPT. Because of the ordering errors the MSTCN cannot be used in standalone mode for assessing segment completion and task completion, since task completion is based on assessment of segment completion and segment order.

The Transformer utilizes raw kinematic features of the torso, upper limb, and object. We experimented with 8, 4, 2 and 1 keypoints from open pose combined with object movement features. We selected the keypoints representing joints that have been show, in our previous work, to have prominence in characterizing upper limb impaired movement (Venkataraman et al., 2014, 2016). It can thus be said that some form of prior expert knowledge is used to constrain the Transformer. The best performing Transformer was with only the wrist key point and object locations. The object location is an important input feature as a lot of the segment transitions are dependent on the object achieving specific translations in the activity space (i.e. movement from bounding box C to bounding box D and back). In prior work, we show that the object-limb end point (wrist or hand) interaction data can be sufficient for the coarse analysis of task completion in upper extremity rehabilitation (Venkataraman et al., 2014). We also further proposed that in these scenarios, the limb could be considered as a dynamic system that can be sufficiently characterized by the behavior of the end point (Venkataraman et al., 2014). Therefore, through the Transformer we codify the long term relationship between the object and the patient limb for better predictions of the segment labels. However, because of the limited and noisy data set, and the limited data points being used for the Transformer, this model has the worst per-frame classification performance of the three segmentation algorithms.

The hidden Markov model (HMM) utilizes kinematic features that are prominent in characterizing upper limb functional movement (Chen et al., 2011b) and also incorporates segment and task layer information through the customized transition matrices. We experimented with multiple training schemes for the HMM. The ablation study of the HMM provides three key insights. First, the HMM is sensitive to input features. The best performance achieved used a six raw kinematic feature HMM model combined with the first derivatives of four kinematic features. The derivatives have more oscillation compared to the raw features as shown in Fig. 8 and follow the Gaussian distribution. Second, the proposed HMM requires 5 transition matrix as a prior. The transition probabilities are selected based on the expert knowledge. The size of the transition matrix depends on the state machine of the exercise. However, we experimented with one transition matrix model to represent all the exercises and as shown in Fig. 7 (b) The performance is poor compared to a 5 model HMM since the expert knowledge of segment-task relation is lost. Lastly, we compared the proposed HMM with a time dependent HMM model but the variance in duration of performance among stroke survivors with different levels of impairment reduces the performance of the time dependent HMM. The HMM has limited segment ordering issues as those are constrained by the transition matrices. However, the reduced sensitivity of the HMM to nuances in the data layer (the HMM model would perform better if the differences in the data between states was significant) combined with the variance in the data cause a higher number of frame level misclassifications. We also varied the batch size to understand the sensitivity of the HMM. In Table 11, the comparative results for different batch sizes are shown. It is evident that with the increase in data size the performance of the HMM increases. Since the HMM tunes its transition matrix and distribution parameters based on the training data, the more data the better the tuning, and we can thus expect to see continuous improvement as we add data in the future.

**Table 11.** Results using different batch sizes for HMM input: (i) Experiment 1: batch size of 1, (ii) Experiment 2: batch size of 9 and (iii) Experiment 3: all data.

|  | Experiment 1 |  | Experiment 2 |  | Experiment 3 |  |
| --- | --- | --- | --- | --- | --- | --- |
|  | Mean | STD | Mean | STD | Mean | STD |
| ACC | 55 | 10.83 | 73.15 | 6.8 | **77.82** | 2.88 |
| Precision | 60.5 | 10.15 | 75.02 | 6.72 | **78.60** | 2.47 |
| Recall | 57 | 10.21 | 75.20 | 5.83 | **78.26** | 2.76 |

**Figure 8.**
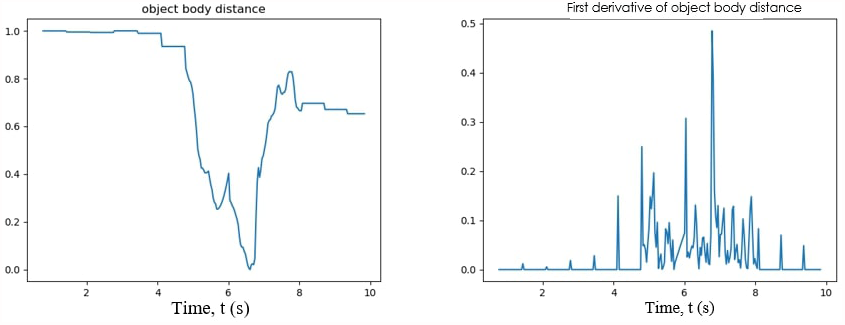
One of the kinematic features representing the distance between the body and the object (left); first derivative of the object body distance feature

**Figure 9.**
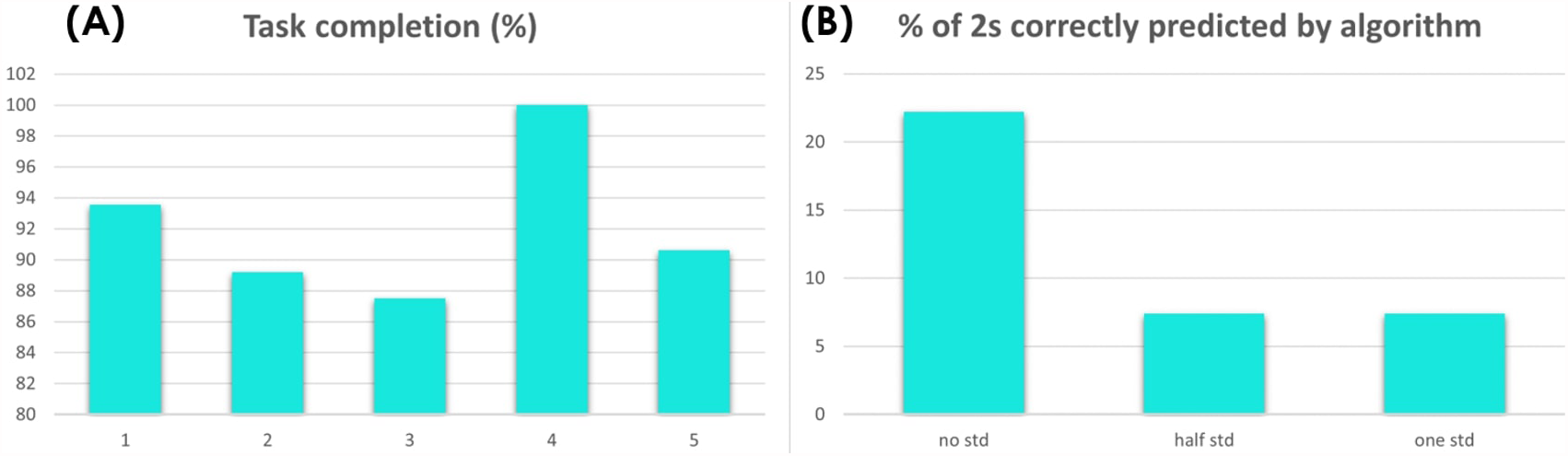
(A) task completion accuracy for 5 splits using the segment blocks (B) percentage of 2s correctly predicted for different range of standard deviation

**Figure 10.**
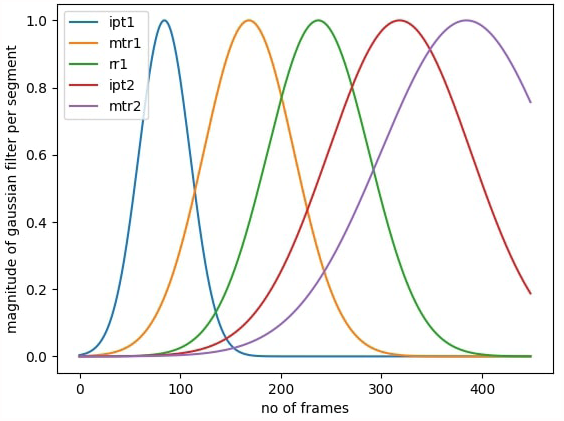
Gaussian windows per segment to filter out false transitions outside the first standard deviation mark

For all the splits, the ensemble model with three algorithms performs the best in the frame level classification. This proves that a fusion of algorithms that have different sensitivities to different layers of the hierarchy outperforms individual models. Without the HMM, the ensemble model predictions have a higher misclassification rate and incorrect sequence orders. For example, without the HMM a prediction can start with MTR or end with IPT. But based on our design prior, we know that all exercises start with IPT and end with *R*&*R*. Since this prior is incorporated into the HMM, the predicted order of segment blocks are correct and the per-frame segment label prediction accuracy increases. As is evident from Table 9, without the MSCTN the ensemble model has significant deterioration in the frame level classification.

To automatically assess the completion of a task, we need to assess the completion and order of segment blocks. This cannot be done solely through the ensemble model as the ensemble model has many false positive blocks and missed transitions. The connection of the segmentation layer to the task assessment layer requires an algorithm that is heavily biased by expert knowledge towards behaviors characterizing transition between segments, as well as the effect of stroke related impairment on these behaviors. The RBBDT algorithm successfully incorporates key unimpaired and impaired movement priors for segment block transition points in one integrated decision sequence. For example, a phase transition is expected for both object and limb end point for a simulated drinking motion (task three) when the object gets close to the head. However, not all stroke survivors are able to lift the object all the way to their mouth, so the phase transition can sometimes happen away from the head. Thus the phase transition becomes the first decision node with the place of the transition in the activity space the second node. The RBBDT is highly sensitive to transition point features but not sensitive to per-frame data patterns across the whole task. Thus the RBBDT produces low quality performance in per-frame classification but higher quality performance in identifying transition candidates. When integrated with the ensemble model, the correct identification of segment blocks rises above 90%.

To perform the task completion assessment, we use a very simple algorithm relying only on comparison of the type and order of the segment blocks given by the integrated ensemble and RBBT predictions, with the expected type and order of segments in tasks established by the expert therapists who designed the SARAH tasks. By adding this codification of prior task knowledge to our analysis, we are able to correctly classify completed and uncompleted tasks over 90% of the time. We are also able to classify completed tasks that took longer than the performance time allowed by the therapist for impaired performance. These results further enhance the validity of the use of a hierarchical model of automated analysis combining algorithms with different sensitivities to the different layers of the hierarchy. This includes layers primarily incorporating observable expert knowledge (higher layers), layers primarily incorporating computer analyzed data (lower layers) and layers integrating data and expert knowledge (middle layers) (see Fig. 1).

The use of approximately 400 videos for training/testing is enough to overcome the issues of overfitting that we demonstrate when using 200 or less videos. As we progressed to 300 and then 400 videos, we show steady improvement for the performance of all the algorithms we were using. It is thus realistic to assume that as we collect more data, the performance of the hierarchical model will continue to improve at all layers.

## 7 CONCLUSION AND FUTURE WORK

Low cost and low intrusion long-term automated rehabilitation at the home is expected to produce low quality and high variability data. A hierarchical model fusing expert knowledge-biased approaches with data-driven techniques can produce robust results in segmentation and task completion assessment in rehabilitation even when utilizing low quality and high variability data. The hierarchical model produces over 90% performance in assessment of segment completion and task completion. Even with this limited information, long term, semi-automated rehabilitation at the home using our SARAH system is feasible. The system can monitor whether the patient has completed the assigned exercises for the day and whether exercises were completed correctly and with ease. The system can use this information to provide coarse feedback to the patient to reward accurate performance or remind the patient of the correct structure of the task if errors are detected. The system can also make simple decisions including when to encourage the patient to repeat an exercise (if they are improving but still have more room for improvement, or move to another simpler or harder exercise (if they have repeated the exercise enough times or are not making progress on the current exercise).

Our computational hierarchy, since it is based on computational data, realizes the hierarchical analysis in a bottom up manner (features conditioning segment frames, conditioning segment blocks, conditioning task completion). Therapists utilize a similar hierarchical analysis in a top down manner (task conditioning segments, conditioning composite features, conditioning raw features) so as to leverage their heuristics about functional task performance. Therefore, the SARAH system can send a daily summary of training results to a remote therapist. Since the computational results will be highly compatible with the therapist assessment heuristics, the therapist will be able to quickly use this summary to structure the next day’s therapy and send a message to the patient guiding their training the next day. The therapist will also be able to select any analysis result and review the corresponding video thus further informing their assessment and therapy adaptation. The feasibility of low-cost semi-automated rehabilitation at the home using the SARAH system will allow the collection of many more videos used to further train our algorithms.

Our team also developed an intuitive assessment interface allowing expert therapists to rate videos of therapy tasks in a top down hierarchical manner, complimenting their assessment approach: rate the overall task, then the segments, then the composite features per segment, and then return to a final assessment of the task). As the videos collected through the SARAH system are rated by expert therapists, we can use this detailed rating (which is highly compatible with our computational approach) to further inform our segmentation and task assessment algorithms and evolve these algorithms to also automatically analyze movement quality and the relation of movement quality to functionality. We are currently planing to use the SARAH system in a pilot study in the homes of stroke survivors in the Spring of 2022.

## AUTHOR CONTRIBUTIONS

There is no special author contribution in this work.

## ACKNOWLEDGMENTS

We thank the therapists of Emory and Carilion hospital for their participation in data capture and rating.

