## Supplementary for "Automated movement assessment in stroke rehabilitation"

### Supplementary Material

#### 1 EVALUATION MATRICES

The accuracy is calculated using,

$$acc = \frac{TP + TN}{TP + TN + FP + FN} \tag{S1}$$

In this formula, TP is the number of positive examples that were correctly labeled, TN is the number of negative examples that were correctly labeled, FN is the number of positive examples that were mislabeled, and FP is the number of negative examples that were mislabeled. To calculate precision/recall we have to sum over rows / columns of the confusion matrix. If the row of the matrix corresponds to specific value for the ”truth”, we can calculate precision/recall using:

$$precision = \frac{M_{ii}}{\sum_j M_{ji}} \tag{S2}$$

$$recall = \frac{M_{ii}}{\sum_j M_{ij}} \tag{S3}$$

That is, precision is the fraction of frames that are correctly labeled as state, $i$ out of all instances where the algorithm labeled $i$. Conversely, recall is the fraction of frames that are correctly labeled as state, $i$ out of all of the cases where the true of state of the frame is $i$.

##### 1.1 Kinematic Features Definitions

###### 1.1.1 End Point and Object Velocity

The end point refers to the wrist coordinates (keypoint 0 in Fig. S1(B)) and $(x_l(t), y_l(t))$ gives us the spatial location of the wrist at time, $t$. We calculate the end point velocity by taking the first derivative of the wrist coordinates w.r.t. time (S4). We use $v_{xl}$ and $v_{yl}$, the magnitude of x and y component of the velocity profile as separate features. We consider the features separately because some of the object transportation in SARAH involve movement in either x or y direction of the image frames.

$$v_{xl}(t) = \frac{d(x_l(t))}{dt}, v_{yl}(t) = \frac{d(y_l(t))}{dt} \tag{S4}$$

Using the bounding box, we calculate a virtual marker that marks the bottom of the object. The x,y-coordinate of this marker represents the location of the object $((x_o(t), y_o(t)))$. The virtual marker enables us to pin point the time frame when the object touches or leaves a base during a transportation segment. Like the end point velocity, we calculate $v_{xo}$ and $v_{yo}$, the x and y component of the velocity profile.

$$V_l(t) = \sqrt{v_{xl}^2(t) + v_{yl}^2(t)}, V_o(t) = \sqrt{v_{xo}^2(t) + v_{yo}^2(t)} \tag{S5}$$

###### 1.1.2 Absolute End Point and Object Velocity

The absolute velocity is the magnitude of the velocity profile given by Eq. S5. The absolute velocity has the signature of starting and ending a movement. Ideally, the magnitude has a bell shaped pattern where it goes from zero to maximum and then decreases to zero again.

$$d_{cl}(t) = \sqrt{(x_c(t) - x_l(t))^2 + (y_c(t) - y_l(t))^2} \tag{S6}$$

$$d_{co}(t) = \sqrt{(x_c(t) - x_o(t))^2 + (y_c(t) - y_o(t))^2} \tag{S7}$$

$$d_{lo}(t) = \sqrt{(x_l(t) - x_o(t))^2 + (y_l(t) - y_o(t))^2} \tag{S8}$$

$$d_{oo}(t) = \sqrt{(x_1(t) - x_2(t))^2 + (y_1(t) - y_2(t))^2} \tag{S9}$$

##### 1.1.3 Relative distances

We calculate 4 distance features that have unique segment signatures for different tasks in the SARAH system. We calculate the distance of the end point and object from the torso line (S6 and S7). The torso line refers to the line joining the shoulder (keypoint 1 Fig. S1(A) and the hip joint (keypoint 8 Fig. S1(A)). We take the center coordinate of this line $(x_c, y_c)$ and calculate distance from it. The distance between the object and end point (S8) signifies the ending and starting time frame of a segment in multiple tasks. Also, calculating distance the between objects (S9) has unique traits of some of the segments that involve object crossing, or putting one object on top of another. We use the first derivative of these distances as input features.

##### 1.1.4 End point Phase Angle

The phase angle is the angle between $v_{xl}$ and $v_{yl}$. The phase angle ($\theta_p$) captures the directionality better than any other features. We calculate the limb phase angle using,

$$\theta_p = tan^{-1}(\frac{v_{yl}}{v_{xl}}) \tag{S10}$$

##### 1.1.5 Relative and absolute end point angle

We calculate two types of end point angle to capture directionality of the end point. The first one is the relative angle between the end point and the torso line. If $b - a$ is the vector joining the wrist and upper torso and $c - b$ is the vector joining the wrist and the lower torso, then the relative angle between the wrist and the torso line is given by Eq. (S11). The second one is the absolute angle or the angular velocity (S12) of the end point that captures the change in directionality in every time steps of the end point movement.

$$\theta_r = Cos^{-1}(\frac{(b-a).(c-b)}{||(b-a)|| * ||(c-b)||}) \tag{S11}$$

$$\theta_a = tan^{-1}(\frac{y_l(t+1) - y_l(t)}{x_l(t+1) - x_l(t)}) \tag{S12}$$

#### 2 RBBDT FEATURE LIST

see Table S1

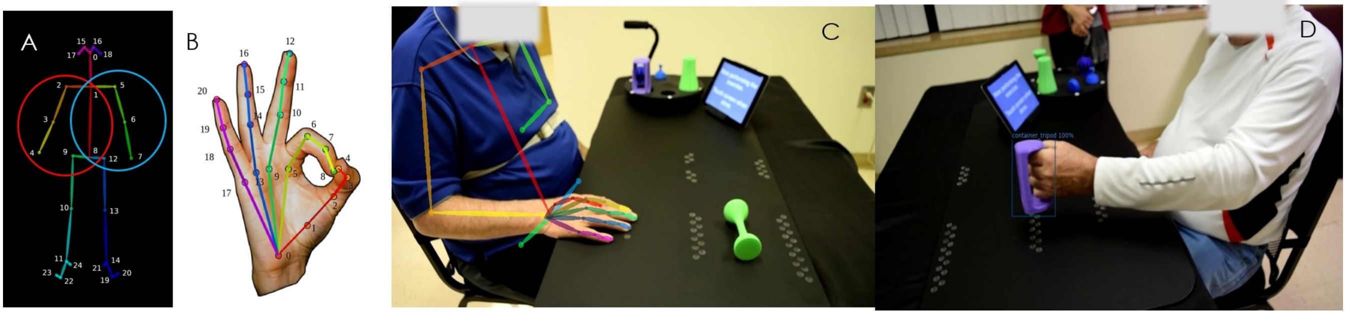


**Figure S1.** (A) 25 body keypoints that OpenPose generates following the COCO dataset; the circles indicate the set of keypoints used in the porposed analysis framework for the right hand imapired patient (red circle) and the left-hand impaired patient (blue circle) (B) 21 hand keypoints (C) OpenPose extracted upper body skeleton overlapped on the actual frame (D) Detected bounding box and object label using Faster RCNN

| Task | Transitions | Features |
|---|---|---|
| 1 and 2 | IPT-R&R | $d_{Fl}, d_{Gl}, d_{Bl}, d_{lo}$ |
| | IPT-R&R | $d_{El}, d_{Fl}, d_{Cl}, d_{Hl}$ |
| | R&R-IPT | $d_{El}, d_{Fl}, d_{Cl}, d_{ol}$ |
| 3 | IPT-MTR1 | $x_o, x_l, d_{Co}, d_{Eo}$ |
| | MTR1-MTR2 | $d_{Eo}, d_{Hl}, d_{Co}$ |
| | MTR2-R&R | $v_{xo}, d_{Hl}, d_{ol}, d_{Co}$ |
| 4 | IPT-MTR1 | $x_o, x_l, d_{Bo}, d_{Ho}$ |
| | MTR1-MTR2 | $d_{Eo}, d_{Hl}, d_{Bo}$ |
| | MTR2-R&R | $v_{xo}, d_{Hl}, d_{ol}, d_{Bo}$ |
| 5 | IPT-MTR1 | $x_o, x_l, d_{Bo}, d_{Ho}$ |
| | MTR1-MTR2 | $d_{Ho}, d_{Hl}, d_{Bo}$ |
| | MTR2-R&R | $v_{xo}, d_{Hl}, d_{ol}, d_{Bo}$ |
| 6 | IPT-MTR | $y_o, y_l, d_{Bo}, d_{Co}$ |
| | MTR-R&R | $V_o, d_{Fl}, d_{Gl}, d_{ol}, \frac{d}{dt}(d_{Co}), \frac{d}{dt}(d_{oo})$ |
| | R&R-IPT | $d_{Fl}, d_{El}, d_{Hl}, d_{Cl}, V_l,$ |
| | IPT-MTR | $y_o, y_l, d_{Bo}, d_{Co}$ |
| | MTR-R&R | $V_o, d_{Fl}, d_{Gl}, d_{ol}, \frac{d}{dt}(d_{Bo})$ |
| 7 | IPT-MTR | $y_o, y_l, d_{Bo}, d_{Co}$ |
| | MTR-R&R | $V_o, d_{Fl}, d_{Gl}, d_{ol}, \frac{d}{dt}(d_{Bo}), \frac{d}{dt}(d_{oo})$ |
| | R&R-IPT | $d_{Fl}, d_{El}, d_{Hl}, d_{Cl}, V_l,$ |
| | IPT-MTR | $y_o, y_l, d_{Bo}, d_{Co}$ |
| | MTR-R&R | $V_o, d_{Fl}, d_{Gl}, d_{ol}, \frac{d}{dt}(d_{Co})$ |
| 8 | IPT-MTR1 | $V_o, x_l, d_{Fo}, d_{Co}$ |
| | MTR1-MTR2 | $d_{Fl}, d_{Gl}, y_l, \frac{d}{dt}(d_{Fo})$ |
| | MTR2-MTR3 | $d_{Co}, d_{Go}, x_l, \frac{d}{dt}(d_{Go})$ |
| | MTR3-R&R | $V_o, d_{Fl}, d_{Gl}, d_{ol}, \frac{d}{dt}(d_{Co})$ |
| 9 | IPT-R&R | $d_{Fl}, d_{Gl}, x_l, d_{Fl}$ |
| | IPT-R&R | $d_{Bl}, d_{ol}, d_{Hl}, V_l$ |
| 10 | IPT-MTR | $y_o, y_l, d_{Bo}, d_{Co}$ |
| | MTR-R&R | $V_o, d_{Fl}, d_{Gl}, d_{ol}, \frac{d}{dt}(d_{Bo}), \frac{d}{dt}(d_{oo})$ |
| | R&R-IPT | $d_{Fl}, d_{Gl}, d_{Hl}, d_{Bl}, V_l,$ |
| | IPT-MTR | $y_o, y_l, d_{Bo}, d_{Co}$ |
| | MTR-R&R | $V_o, d_{Fl}, d_{Gl}, d_{ol}, \frac{d}{dt}(d_{Co})$ |

**Table S1.** List of RBBDT Features used for different segment transitions in different exercise
